# Establishment and characterisation of oviductal organoids from farm and companion animals

**DOI:** 10.1101/2022.11.05.515300

**Authors:** Edwina F. Lawson, Arnab Ghosh, Victoria Blanch, Christopher G. Grupen, R John Aitken, Rebecca Lim, Hannah R. Drury, Mark A. Baker, Zamira Gibb, Pradeep S. Tanwar

## Abstract

Organoid technology has provided us with a unique opportunity to study early human development and decipher various steps involved in the pathogenesis of human diseases. The technology is already used in clinics to improve human patient outcomes. However, limited knowledge of the methodologies required to establish organoid culture systems in domestic animals has slowed the advancement and application of organoid technology in veterinary medicine. Here, we have developed a platform to grow organoids from animal tissue samples and characterized oviductal organoids from five domestic animal species. Organoids were grown progressively from single cells derived from the enzymatic digestion of freshly collected equine, bovine, feline, canine, and porcine oviducts. The addition of WNT, TGFB, BMP, Rock, and Notch signalling pathway activators or inhibitors in the culture medium suggested remarkable conservation of the molecular signals involved in oviductal epithelial development and differentiation across species. The gross morphology of organoids from all the domestic species was initially similar. However, some differences in size, complexity, and growth rate were observed and described. Well-defined and synchronised motile ciliated cells were observed in differentiated organoids in mature populations. Histopathologically, oviductal organoids mimicked their respective native tissue. In summary, we have developed a detailed cross-species comparison of oviductal organoid models, which will be valuable for advancing assisted reproductive technologies and fertility studies in these animal species in the future.

**Summary sentence:** Organoids can be derived from the oviductal epithelium of cow, cat, dog, horse, and pig to advance assisted reproductive technologies in animals.

## INTRODUCTION

The oviduct is not solely a conduit connecting the ovary to the uterus, but a complex hormonally regulated organ, which is essential to sustain fertilisation and early embryonic life (Lyons et al., 2006). The oviduct, known as the fallopian tube in humans, consists of four functionally unique segments. Most proximal to the ovary are the fimbrial branches of the infundibulum, which pick up and propel the newly ovulated oocyte to the ampulla. The ampullary region is where spermatozoa undergo the final stages of capacitation and achieve fertilisation of the oocyte. This region is connected to the isthmus, which is believed to play roles in both the storage of spermatozoa prior to ovulation and the regulated transport of cleaving embryos. Lastly is the proximately located utero-tubal junction, (Talbot et al., 1999), which is simultaneously a major barrier for sperm transport (Suarez and Pacey, 2006), and the final region embryos await before reaching the uterine lumen (Avilés et al., 2015). The lumen of the mammalian oviduct is lined by intricately folded pseudostratified epithelial cells, containing interspersed secretory and ciliated cells (Ito et al., 2020). Ciliated cells propel the oocyte by creating luminal currents through continual synchronised beating of their cilia axoneme (Ezzati et al., 2014), whereas the secretory cells produce the complex oviductal fluid, which provides the optimal micro-environment for sperm capacitation, fertilisation and early embryonic survival (Abe and Oikawa, 1993, Chen et al., 2013).

Like many epithelia, the oviduct contains a population of stem cell-like progenitors, which actively divide to facilitate epithelial regeneration and homeostasis. Oviductal fimbriae epithelial cells are regularly exposed to follicular fluid during each ovulation (Huang et al., 2015). The follicular fluid contains copious inflammatory cytokines and reactive oxygen species (ROS) which readily lead to tissue injury and DNA strand breakage in the cells of this region (Hamdan et al., 2016, Al-Saleh et al., 2021). Thus, the oviductal fimbria epithelium is one of the tissues with most abundant stemness activity, meaning it is required to maintain a balance between quiescence, proliferation, and regeneration. As such, in the epithelium, secretory cells are the most frequently dividing cells, which not only self-renew, but also give rise to ciliated cells. Lineage tracing work has identified that such stem-cells express the secretory marker PAX8 (Ghosh et al., 2017), which differentiate into ciliated cells (acetylated a-tubulin positive) through the WNT-signalling pathway and play a crucial role in oviduct development and homeostasis (Kessler et al., 2015). Furthermore, most proliferating cells in the oviductal epithelium are secretory cells (Ghosh et al., 2017), which in humans and the mouse may be driven to differentiate into fallopian/oviductal organoids when supplemented with niche factors (Kessler et al., 2015, Xie et al., 2018), further supporting their role as stem/progenitor cells. Single cell sequencing analyses of human fallopian tube epithelial cells have identified several different populations of these secretory cells and further studies are required to clarify their contributions to oviductal epithelial homeostasis (Ahmed and Tait, 2020, Dinh et al., 2021).

Recent advances in assisted reproductive technology (ART) have helped us to better understand the causes of infertility. Even though there have been great developments in ART and the importance of the oviduct for optimal gamete maturation, capacitation, and embryo development has been established (Avilés et al., 2010), many unanswered biophysical questions remain for domestic species. For example, in the equine, the biochemical signal for maternal recognition of pregnancy (MRP) has not yet been identified (Swegen, 2021) and an efficient *in vitro* fertilisation (IVF) protocol has not yet been established (Leemans et al., 2016). In cattle, IVF is widely used in commercial settings but *in vitro* conditions for fertilisation and embryo production are still far from optimal. As such, it is believed that up to 50% of pregnancy failures in dairy cattle take place in the oviduct (Walsh et al., 2011). Furthermore, polyspermy is a frequent phenomenon during IVF in many species, including porcine, while in an *in vivo* setting it is rarely observed (Lawson et al., 2022). For companion species, such as the canine and feline, IVF is not well established, and a detailed understanding of reproductive idiosyncrasies has predominantly been established for the purpose of application to endangered species of non-domestic taxon (Thongphakdee et al., 2020). These gaps led to the hypothesis that a species-specific oviductal organoid model could be uniquely developed, and characterised for individual species. Oviductal studies have the potential to advance IVF in domestic species by allowing detailed *in-vitro* studies to be carried out on the microenvironment in which fertilisation and early embryo development occur. Domestic animal research has the potential to improve food security and welfare outcomes, while providing models for human health research (Seeger, 2020). This latter contribution is recognised by the One Heath Initiative, in which the close linkage between the health of humans, animals and the environment is widely accepted (Seeger, 2020). As such, organoids will be valuable for evaluating cross-species disease variations or species-specific infectivity of zoonotic pathogens, while providing an *in vitro* alternative for infection studies (Kawasaki et al., 2022).

In a clinical setting, organoids may be cultured from specific patients to conduct *in vitro* drug efficacy testing for tailored treatments (Berkers et al., 2019), facilitating personalised medicine into clinical practice. Oviductal organoids composed of fully differentiated ciliated and secretory epithelial cells have been successfully generated from human (Kessler et al., 2015, Chang et al., 2020) and mouse-derived cells (Xie et al., 2018, Ghosh et al., 2020). While these systems have vastly improved the understanding of pathologies and the process of fertilisation in each of these species, their applicability to other animals is limited because oviductal physiology and fertilisation strategies vary greatly between mammalian species. As such, the aim of this study was to carry out a detailed cross-species comparison of oviductal organoid development, with the intention to provide advancements for both ART and studies of fertility.

## MATERIAL AND METHODS

### Oviduct epithelium isolation and organoid culture

Whole oviducts were obtained from adult porcine (n=3), bovine (n=2), equine (n=4), canine (n=5) and feline (n=4) females, and samples were collected either directly from the slaughterhouse or abattoir floor or collected from discarded post-surgical hysterectomy material. As this work was conducted using opportunistic sourcing of discarded tissues and organs, no formal institutional animal ethics committee approval was required. After collection, samples were then washed and transported on ice in a transportation medium (Dulbecco’s Modified Eagle Medium/Nutrient Mixture F-12 (DMEM-F12 cat # 12634-010) with 1% penicillin-streptomycin (Life Technologies, cat # 15070-063)) and stored in at 4°C overnight in the same medium. Oviducts were washed three times with Dulbecco’s phosphate-buffered saline (DPBS) and 1% penicillin-streptomycin to remove the blood and unwanted debris. Multiple small portions of the fimbriae were dissected under a stereomicroscope and digested in Accumax (Innovative Cell Technologies, Inc., cat#AM105) enzyme for 2-3 hours on a shaker at room temperature. Enzymatic digestion was stopped by adding a complete medium comprising 5% foetal bovine serum (FBS), 1% Glutamax (Sigma), 1% HEPES and 1% penicillin-streptomycin in Advanced DMEM-F12. The cell suspension was collected in a Falcon tube and centrifuged at 1500 rpm for 5 minutes to collect the epithelial cells. These epithelial cells were then propagated in a complete medium supplemented with the addition of human Epidermal Growth Factor (hEGF) (12ng/ml, PeproTech), ROCK inhibitor (Y-27632; 10 mM; TOCRIS) and Primocin (0.2%) as a 2-dimensional culture on a matrigel coated cell culture plate. Once 70% confluence had been achieved, epithelial cells were detached by trypsinisation and centrifuged at 1500 rpm for 5 min to collect the epithelial cells. Epithelial cells in a fresh complete medium were then incubated for 2.5-3 h at 37°C, in a 5% CO2 incubator for differential attachment to enrich pure epithelial cells. After epithelial cell enrichment, the cells were counted and resuspended in a complete medium without FBS, referred to as Oviductal Epithelial (OE) culture medium. Following this, the epithelial cells were mixed with matrigel and placed as a 50 μl drop in each well of a 24-well cell culture plate, with 25,000 cells in each 50 ul drop. The matrigel with cells was allowed to solidify by keeping the 24-well plate at 37°C for 20 minutes. Then each well of the culture plate was overlaid with 75% OE medium and 25% WNT3A-RSPO3-NOGGIN conditioned medium (WRN-CM) prepared from L-WRN cells by following a previously described protocol (Miyoshi et al., 2013).

The medium was further supplemented with growth factors and signalling modulators comprising of human Epidermal Growth Factor (hEGF) (12ng/ml, PeproTech), B27 Supplement (2%, Gibco), ROCK inhibitor (Y-27632 dihydrochloride; 5 mM; TOCRIS), N-Acetyl-L-Cysteine (1.25 mM/ml, Sigma), Nicotinamide (10 mM/ml, Sigma), Primocin (0.2%, Invivogen), and A83-01 (0.5 mM, TOCRIS). The medium for organoid culture was replenished every three days and on day 21 the organoids were harvested for further processing. For each sample, 25-30 organoids were considered at each time point for each species.

### Immunofluorescence (IF) and Hematoxylin & Eosin (H&E)

Oviductal tissues were fixed in 4% (w/v) paraformaldehyde at 4°C overnight before being processed for paraffin embedding. Organoids were fixed in 4% (w/v) paraformaldehyde for one hour at room temperature. The paraffin-embedded blocks were then sectioned at 5 μm thickness. For IF and H&E staining, 5 μm sections were deparaffinised followed by rehydration, and processed separately according to protocol. For immunofluorescence staining, antigen retrieval was carried out on rehydrated sections by heating them to either at 110°C (oviductal tissue) or 98°C (organoids) for 30 minutes in EDTA buffer (1mM; pH 8.0). After blocking, sections were incubated with primary antibodies at 4°C overnight. The primary antibodies used were rabbit anti-PAX8 (1:1000; catalog number 10336-1-AP; Proteintech), mouse anti-Acetylated-Tubulin (1:1000; catalog number T7451, clone 6-11B-1, Sigma-Aldrich), rabbit anti-Ki67 (1:400; catalog number Ab15580, Abcam) and DAPI dilactate (Sigma-Aldrich D9564). The secondary antibodies were Alexa 488/594 conjugated anti-rabbit/mouse IgG (1:250; Jackson ImmunoResearch Labs).

### Microscopy and image acquisition

Initial gross oviductal images were captured using Nikon SMZ25 stereoscope. During organoid culture, organoid development images were taken by JuLiTM Stage Real-Time Cell History Recorder (NanoEnTek) every 3 days. Fluorescence and H&E images were acquired using Olympus DP80 CCD (charge-coupled device) and cellSens software (Olympus). Confocal laser scanning was carried out on a Zeiss LSM 900-Airyscan-2 microscope with a 63x objective using the software ZEN (blue edition) 3.0 and were reconstructed by sequences (z-stack), organoid imaging measurement and analysis software was carried out using Fiji 9 (ImageJ) (Schindelin et al., 2012).

Mature organoids containing cilia were transferred to a recording chamber at room temperature (22–25 °C), containing OE culture medium and perfused at a rate of two bath volumes per minute (i.e. 3 ml/min), Cilia was observed under a Zeiss Axioskop microscope using infra-red differential interference contrast optics and a Zeiss AxioCam Mrm camera. From the equine and porcine organoid populations, cilia were identified and imaged using Zen Blue software (version 2.6) and live ciliary motions were recorded from 12 regions of ciliated axonemes at 30 second intervals. The recording captured a synchronous and continual beating of motile cilia which generated a coordinated and unidirectional flow in both species (Supp. 1).

## RESULTS

### Gross and histological characterization of oviducts from domestic animals

Healthy feline, canine, porcine, bovine, and equine oviducts were collected, and gross images of morphological infundibular characteristics of each species were obtained, as shown (Figure 1i). Despite the infundibulum serving the same function across the species, namely, to collect ovulated oocytes and transport them, clear morphological differences were evident. Macroscopically, the infundibulum presents with funnel-shaped irregular projections (white arrows) with branching fimbriae engulfing the ovary. Interestingly, cat and dog infundibulum adhere tightly to the ovary (Fig1i A, Fig1i B). In the case of the porcine, equine, and canine, the fimbriae fonds branch more loosely around the ovary without being attached (Fig1i C, Fig1i D, Fig1i E,). Oviductal sectioning, followed by H&E staining (Figure 1ii) showed the histological architecture of native oviducts, including all three distinct layers, serosa, muscularis and mucosa. The oviductal epithelium from each species comprised a single convoluted layer of columnar epithelial cells lining the lumen, creating both luminal projections and differently shaped crypts.

**Figure 1.**
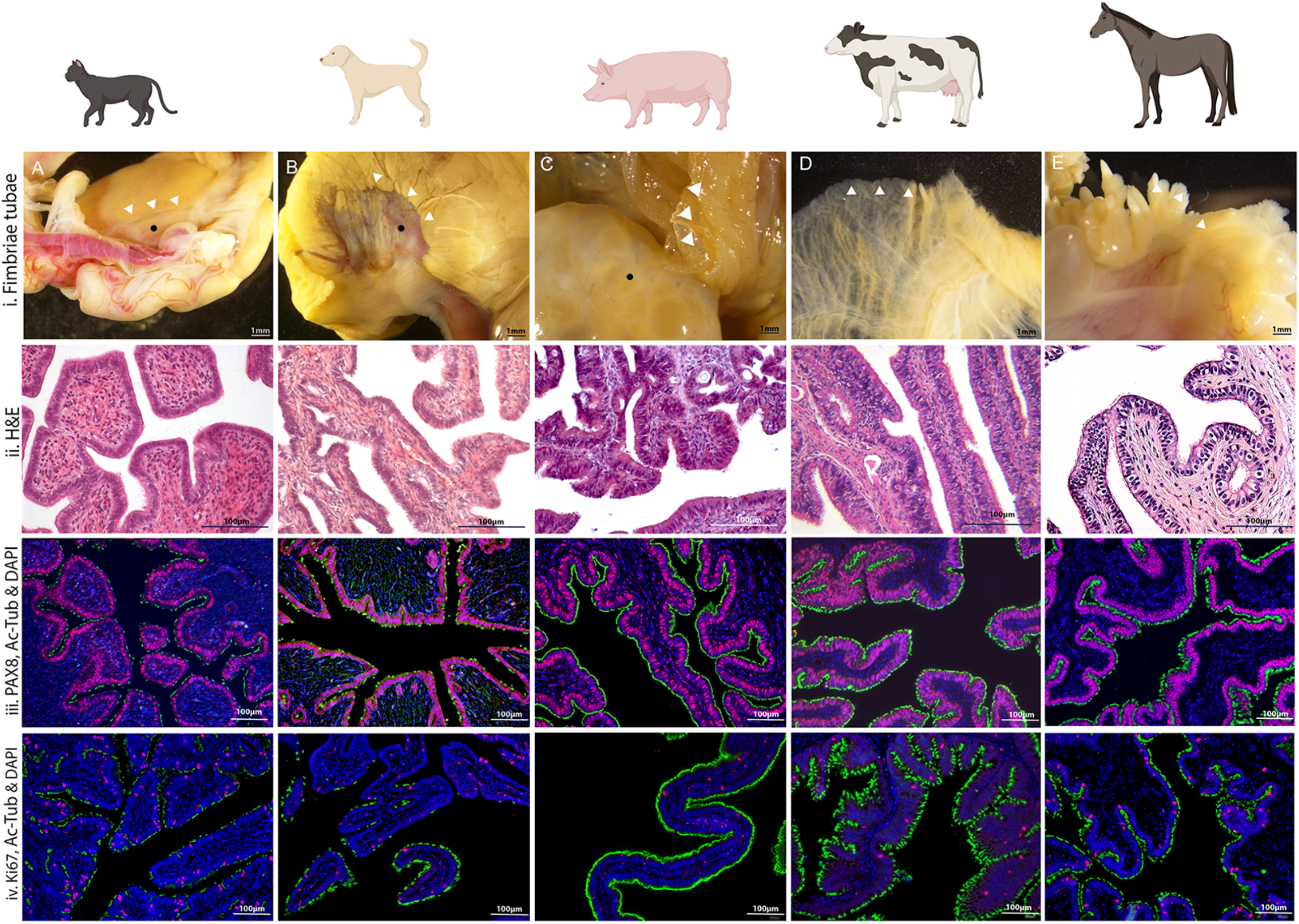
Characterisation of feline (A), canine (B), porcine (C), bovine (D) and equine (E) native oviductal tissue. i) Gross anatomical features of oviductal fimbriae including the ovary in feline, canine, and porcine images (black circle). The fimbriae of the infundibulum branches to tightly engulf the ovary (white arrow) in feline and canine. In the porcine, bovine and equine, the fimbriae projections are not adhered to the ovary but freely interact with the ovarian structure. ii) Histological staining with H&E details the infundibular epithelium, lined with a single layer of polarised columnar epithelial cells. In each species, a highly folded mucosa with crypts is evident. iii) Immunofluorescence imaging showing a consistent expression of PAX8 (red) positive secretory epithelial cells, located mostly along the basal region of the epithelial cells. Additionally, Ac-tubulin (green) is depicted staining the flagella-like projections of the epithelial ciliated cells. In the cat and dog Ac-tubulin positive ciliated cells were more irregularly spaced, compared to that in porcine, bovine, and equine. DAPI (blue) staining shows the nuclei of cells. iii) Immunostaining depicting positive staining for Ki67 (red), strictly associated with cell proliferation constitutively. Cells can be seen dispersed throughout normal healthy oviductal epithelium. Additionally, mature ciliated cells are stained positive for Ac-tubulin (green) and cell nuclei are stained with DAPI (blue).

### Distribution of secretory and ciliated cells in oviducts

Immunofluorescence (IF) analysis of native oviductal epithelium showed a consistent expression of PAX8-positive secretory epithelial cells located mostly along the basal region with a cohesive apical-to-basal polarity (Image 1iii). Using DAPI as a nuclear counterstain, acetylated α-tubulin (Ac-Tub) was used to identify the flagella-like projections of the ciliated cells by staining the acetylated microtubules found within centrioles and cilia. Notably, in the cat and dog, Ac-Tub positive ciliated cells were consistently expressed but punctate and more irregularly spaced throughout the infundibulum (Fig1iii A, Fig1iii B), compared to porcine, bovine, and equine oviducts (Fig1iii C, Fig1iii D, Fig1iii E). Additionally, Ki67, the nuclear DNA-binding protein strictly associated with cell proliferation, was constitutively dispersed throughout normal healthy oviductal epithelium (Fig 1iv). The presence of Ki67 and its distribution suggests that cells are actively dividing.

### Establishment of oviductal organoids cultures

The fimbrial end of equine, porcine, bovine, canine, and feline oviducts were collected. Single cells were isolated using enzymatic digestion. These single cells were embedded in matrigel and supplemented in the culture medium with required growth and niche factors. Organoids progressively developed from single cells over the course of 21days. The organoids were imaged every 3 days (Fig. 2), to record a timeline of the visual change occurring in the cultures, which included recording morphological changes and size, as measured by perimeter (μm). During the initial phase of growth, no ciliated cells or ciliary beating were observed, however, these cells appeared after 10 days of cultures. Cultures were efficiently generated long-term and maintained for several months. Organoids were also successfully grown from cells frozen in liquid nitrogen.

**Figure 2.**
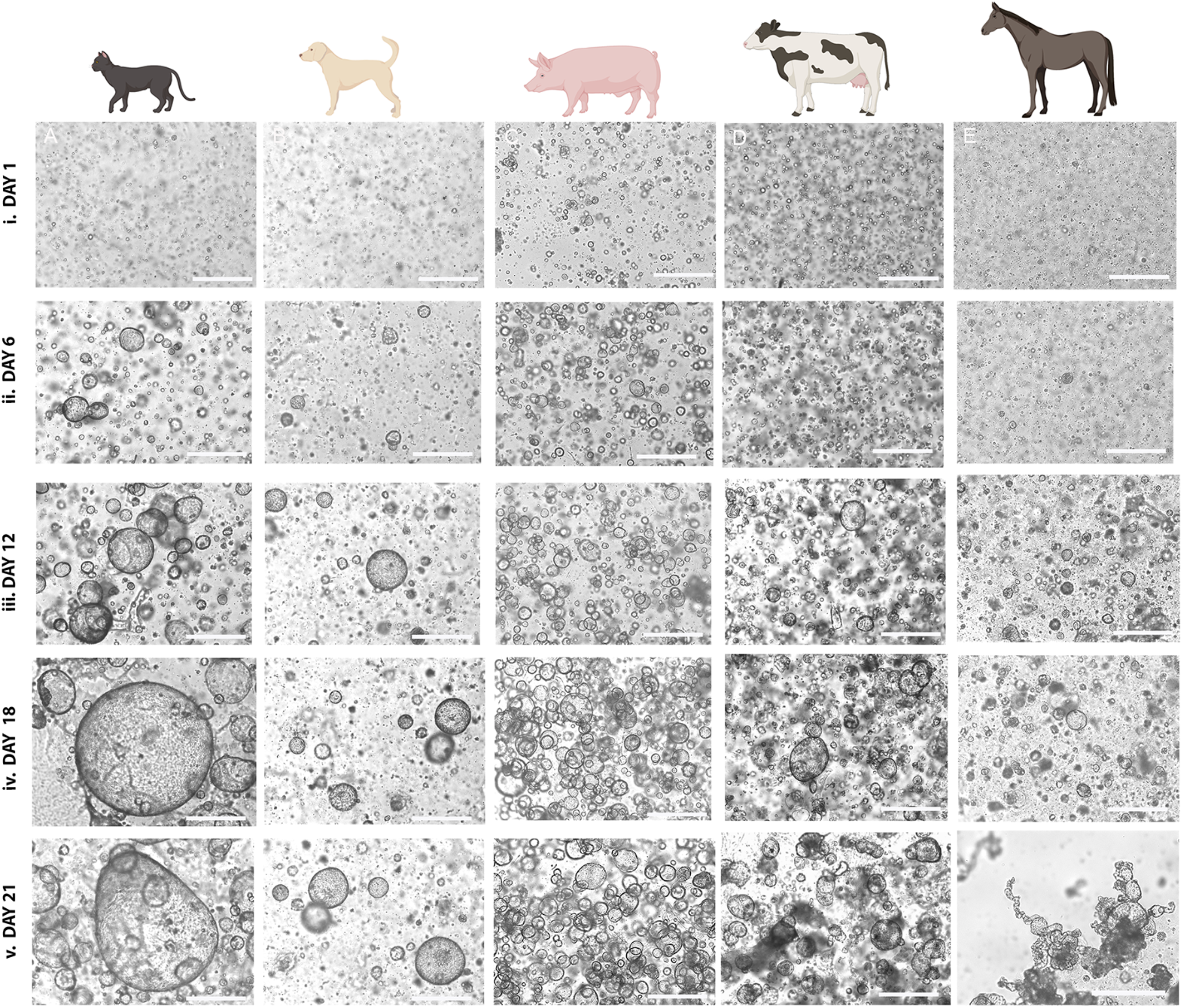
Representative bright-field microscopy images showing organoid development from feline (A), canine (B), porcine (C), bovine (D) and equine (E) oviductal epithelial cells at Days 1, 6, 12, 18 and 21 of culture (I, ii, iii, iv and v, respectively). Maximal individual organoid expansion can be observed occurring between Days 12 and 18. The scale bars shown on all images correspond to a length of 500 μm.

### Species variation in organoid phenotype and growth

For each species, cultured organoid growth was consistent, but between species, organoid expansion and size progressed along different trajectories. From day 6, a marked expansion in size was observed for each species (Fig2ii). However, feline oviductal organoids had exponential growth in size (perimeter) from day 6 (Fig2ii.A), showing a notably higher rate of expansion from that point compared to other species (Fig2ii B,C,D,E). Collectively, maximal organoid size growth occurred between day 12 and day 18 (Fig2iii). In addition to sharing prominent morphological characteristics, such as ciliation, phenotypic differences were immediately apparent between the species. Evident after day 18, equine and bovine organoids lost the spheroid phenotype (Fig2iv.D, E), and by day 21 more complex cell configurations developed. Organoids from all species developed crypts and villi-like structures and internal folding architecture and invaginations, which were analogous to the mucosal structure of the native oviduct (Fig2v.D, E). The branching architecture was particularly and consistently evident in the equine, as highlighted by comparing the morphological differences of mature feline (Fig 3A) and equine oviductal organoids (Fig 3B). To determine if the organoid cell-network architecture was similar to that observed in the native oviductal epithelium (Fig 4), H&E staining was carried out on oviductal organoid full-thickness sections (Fig 4i) after 21 days of culture. Staining showed that oviductal organoids were composed of a monolayer of polarised columnar epithelial cells, exhibiting equivalent complex internal folding architectural structures.

**Figure 3.**
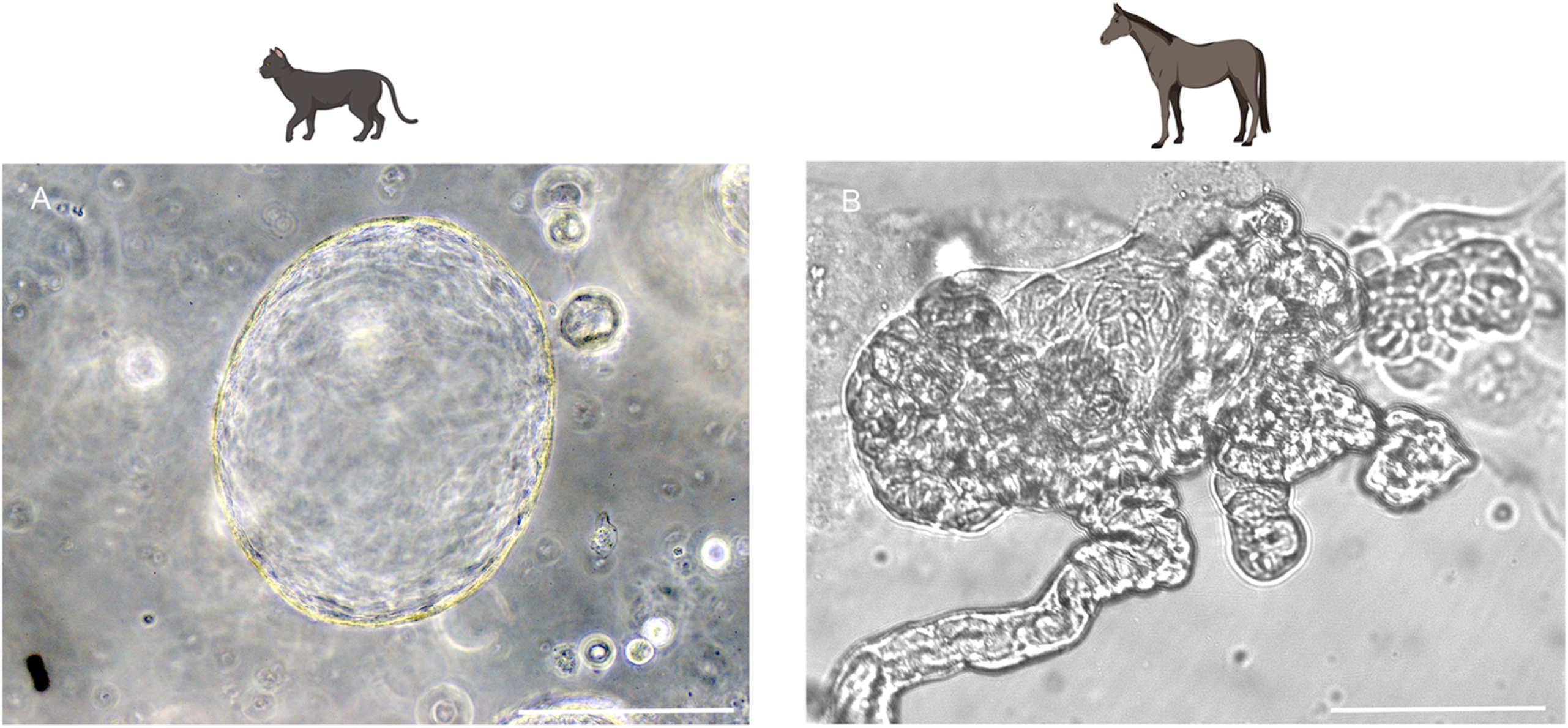
High magnification (x20) brightfield images of feline (A) and equine (B) mature organoids at 21-days of culture. The equine organoids started branching as they matured and depicted a more varying and complex shape compared to the feline oviductal epithelial organoids, which remained more spheroid in structure. The scale bars correspond to a length of 125 μm.

**Figure 4.**
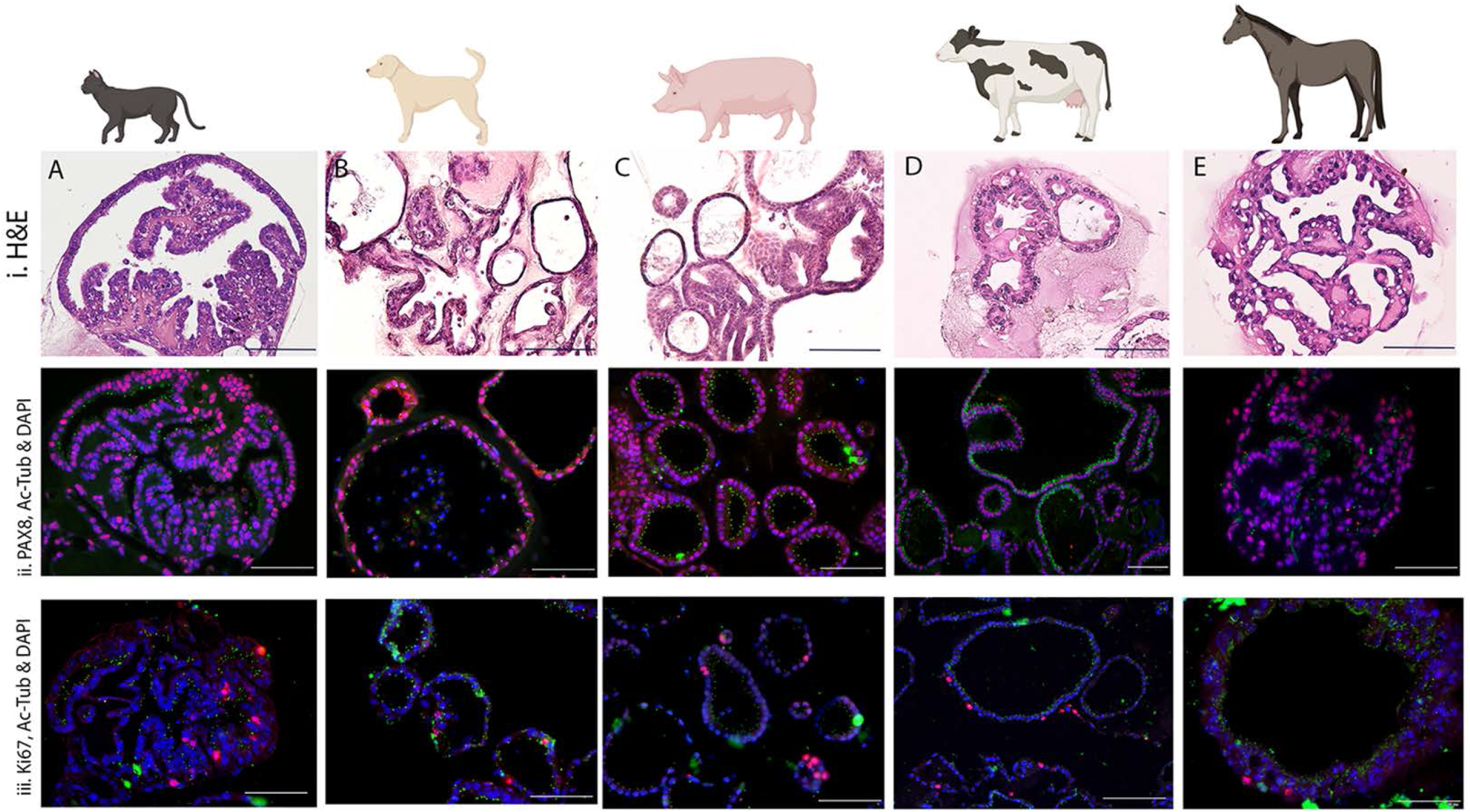
Characterisation of feline (A), canine (B), porcine (C), bovine (D) and equine (E) oviductal epithelial organoids at 21 days of culture. i) Histological staining with H&E showing oviductal epithelial organoids with a monolayer of polarised columnar epithelial cells lining the lumen. In each species, a clear lumen is evident characterised by the presence of luminal crypts and folds. ii) Immunostaining with PAX8 (red), Ac-Tubulin (green), and DAPI (blue) within epithelial cells. Positive staining for α-tubulin indicates ciliated cell differentiation. Positive PAX8 nuclear staining indicates the presence of progenitor secretory cells, which are non-ciliated. DAPI effectively stained the nuclei. Staining patterns mimicked that represented in native oviduct tissue. iii) Positive identification of the marker for proliferating cells, Ki67 (red), showed it was constitutively dispersed and basally located throughout. Positive nuclear staining for Ki67 suggests that the cells were actively dividing. In addition, Ac-Tubulin (green) stained ciliated cells were identified together with the DAPI (blue) stained nuclei of cells. The scale bars shown on all images correspond to a length of 100 μm.

### Immunostaining of organoid cultures

For further comparison, individual cultures were stained with PAX8, Ac-Tub and Ki67, with DAPI serving as a nuclear counterstain after 21 days of culture (Fig 4ii). Mature organoids consistently expressed PAX8-positive secretory cells as well as PAX8-negative Ac-Tub-positive ciliated cells (Fig 4iii). Given that the highly proliferating PAX8 cells respond to clear Wnt signalling, which modulates differentiation into ciliated cells (Stewart and Behringer, 2012), their presence in oviductal organoids is indicative of epithelial cell homeostasis. Interspersed Ki67-positive proliferating cells were detected scattered at the basal layer of the epithelium (Fig 4iii), demonstrating active proliferation in each population. Interestingly, the IF revealed that much of the expression patterns in oviductal organoids, such as the proliferation patterns of Ki67 positive cells, and the consistency of PAX8, Ac-Tub positive cells, consistently mimicked native oviductal epithelium.

### Functional analysis of cilia

Acetylated alpha-tubulin-positive cilia were identified, projecting luminally in mature organoids of all species. Moreover, across all species, and consistent with native oviductal epithelium. The presence and detail of Ac-Tub-positive cells with visible cilia were captured with high magnification using IF confocal imaging (Fig 5), such ciliated cells contained abundant well-defined cilia that were apically pointed and anchored to well-aligned basal bodies. For functional validation of the model, mature organoids should possess a functional ciliary axoneme capable of generating a unidirectional flow and a ciliary beat. As such, motile cilia were observed consistently in the mature organoid population, with mature cilia vigorously motile and recorded using high-speed infra-red differential interference contrast optics. A synchronised, continual beat was recorded in both equine and porcine with actively motile cilia generating a coordinated and continual, unidirectional flow (Supp. 1), of secretion, debris, and medium.

**Figure 5.**
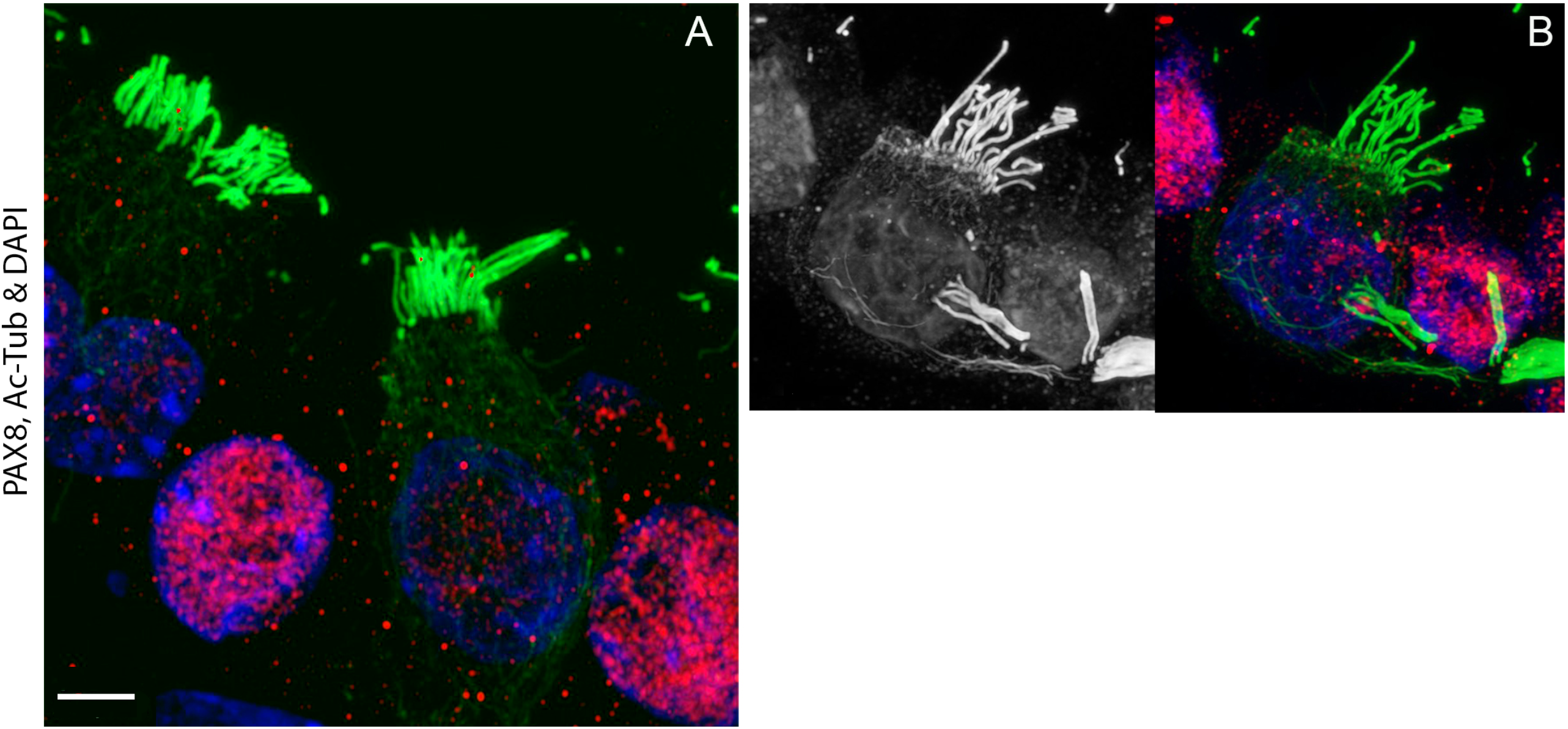
Representative immunofluorescent confocal images showing detailed cilia structures of porcine organoids at 21-days of culture. Ciliated cells contained abundant well-defined cilia, predominantly apically pointed, and anchored to well-aligned basal bodies. Those cells with acetylated microtubules showed positive staining (green) with Ac-Tubulin but were negative for PAX8 secretory cells and vice versa (A). In some of these cells, Ac-Tubulin positive cilia can be observed projecting into a different confocal plane of view (B). The scale bars correspond to a length of 4 μm.

**Figure 6.**
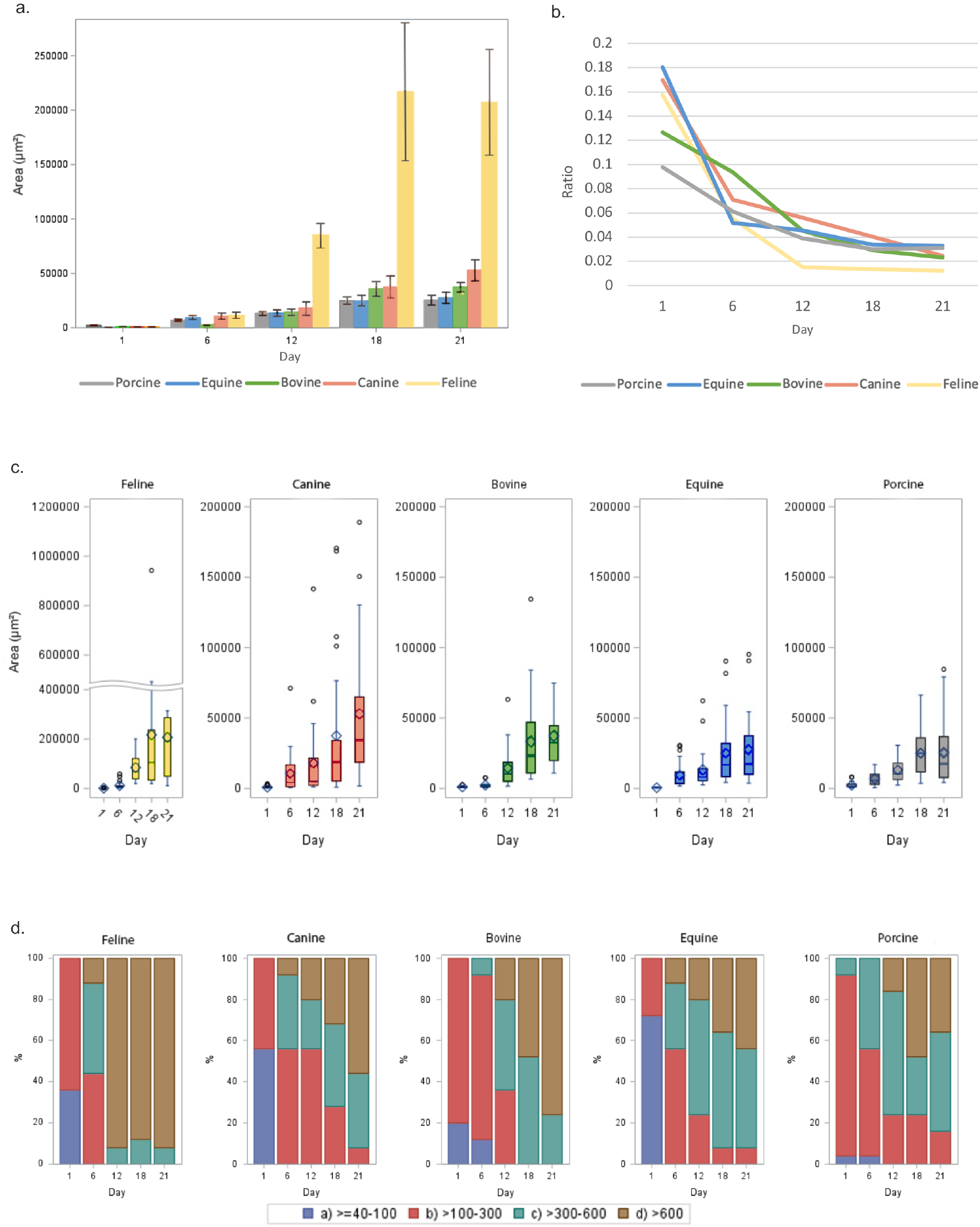
Quantification of organoid development from feline, canine, porcine, bovine and equine oviductal epithelial cells, over a period of 21 days in culture. a) Quantification of the average area (μm^2^) organoid size between species, at 1, 6, 12, 18 and 21 days of culture. b) Each species represented via complexity index. Perimeter to area ratios measure the complexity of the 2D biological shape between organoid species. The higher the value the more complex the shape via the application of morphometrics. c) Quantification and comparison of organoid sizes for each species, with area growth per day. d) The distribution of organoid sizes over 21 days of culture according to area (μm^2^).

### Quantification of organoid development

For quantification of organoid development, individual organoids (n=25) were measured from each sample, for each species, and the mean area (μm^2^) recorded, at 1, 6, 12, 18 and 21 days of culture (Fig6a). To quantify the distinct changes in morphology, the principles of morphometrics were applied using perimeter to area ratio to measure structural complexity (Gardiner et al., 2018, Posamentier and Lehmann, 2011). Canine, porcine, bovine, and equine organoids displayed a high measure of biological structure complexity (Fig6.B), particularly when compared to the rapidly enlarging perimeter observed in the mature feline organoids. Importantly, the oviductal organoid populations developed from single cells and progressively grew in size. For each species, this size was compared, using area (μm2) growth per day (Fig6C) and the distribution of organoid sizes over 21 days of culture was recorded (Fig6D).

## DISCUSSION

The purpose of this study was to develop and characterise adult stem cell-derived oviductal organoid cultures across multiple domestic species. By developing a standardised model, this study found that the mature oviductal organoids that were produced resembled their source tissues architecturally, structurally, and functionally. In this study, a standardised model was successfully generated. Initially, internal architecture was shown to develop in individual organoids, and mimic the internal architecture of native organoidal epithelia. Such architecture was highlighted using H&E staining and reflected native oviductal tissue (Fig 1). It has previously been shown that within oviductal epithelia, the secretory cells continuously replenish, by transitioning to ciliated cells (Ghosh et al., 2017). In the present study, the progenitor secretory cell marker PAX8 was consistently present in mature oviductal organoids in all species investigated (Fig 4i). The presence of such progenitor cells suggests that oviductal regeneration and homeostasis is occurring in the organoid populations. Furthermore, Ki67, a marker of proliferation, was also observed in organoid populations, indicating that they were actively dividing (Fig 4ii). The identification of Ki67 and PAX8 in these organoids is an important finding, but even more notable was the presence of mature, functionally motile ciliated axonemes in organoid populations in 21 day organoids. Cells containing mature ciliated axonemes were initially identified by the presence of Ac-Tub (Fig 5), and detailed images of these structures were captured. In addition, these cilia were observed to produce a synchronous and continual beat frequency resulting in a unidirectional flow (Supp 1). As the mammalian oviduct is one of the organs that relies heavily on the motility of cilia for its normal functions (Wang et al., 2015), the identification of actively motile cilia supports the functionality of the organoid model described in this manuscript.

Anatomically, the proportion of ciliated cells is most numerous in the distal oviduct (fimbria, infundibulum and ampulla) (Stewart and Behringer, 2012). Previous studies have shown that stem-like cells are found in the fimbrial region of the oviduct in the mouse and human (Das et al., 2006, Yamamoto et al., 2016, Paik et al., 2012, Patterson and Pru, 2013), and include PAX8 progenitors cells, that transition, giving rise to ciliated cells (Ghosh et al., 2017). Given that fimbrial cells possess the highest organoid-forming capacity (Xie et al., 2018, Rose et al., 2020), it was this anatomical region which was harvested for this study. However, the collection of such oviductal specimens, was limited to those discarded samples, as such, the cyclicity, precise age and exact health status could not always be accounted for in the present study. Yet, the findings seem to provide a framework for models in species other than human or mouse (Rose et al., 2020, Kessler et al., 2015).

Although mature organoids were observed in all species by 21 days, the interspecies differences in oviductal development rate and structure was not surprising. Many mammalian species display unique features in fertilisation physiology and early embryo development. For example, in the bitch, the oocyte is released two or three days prior to fertilisation (Avilés et al., 2015), rather than at the time of ovulation. In the mare, the developing embryo remains in the oviduct for 5 days after fertilisation (Betteridge et al., 1982, Freeman et al., 1991), contrasting markedly with the cow and sow, in which the embryo/s remain in the oviduct for only 3 and 2 days respectively (Hafez and Hafez, 2016, Dziuk, 1985). In addition, parity size should be taken into account when considering the function of the oviduct; the sow can carry 16 piglets per gestation, whereas in the mare, similar to the human female, only one offspring is typically carried to term. Such functional idiosyncrasies must be considered when comparing oviductal epithelium. In this study, the comparative differences in organoid size (seen in the feline) and complexity observed (in the equine) may indeed relate to the different patterns of early embryonic development between species. The different growth trajectories and proliferation rates of the organoids at different time points during *in vitro* culture may reflect the species-specific differences in oviductal tube physiology (Avilés et al., 2010). Given that reproductive science, plays a critical role in human health, it is remarkable that approximately only 10% of mammals have been studied in this field (Wildt et al., 2010). Knowledge gained from such cross-species comparisons highlights the diversity that exists (Nagashima and Songsasen, 2021), and how the lack of species-specific studies in reproductive physiology contributes to reduced fertility rates. In a commercial setting, poor fertility often translates to financial losses for producers and compromised welfare arising from repeated invasive veterinary interventions (Li and Winuthayanon, 2017). So, although IVF techniques have progressed over the last 40 years, and are now used in multiple domestic and non-domestic species around the world (Sirard, 2018), they are not completely matured; and a number of problems remain unsolved or not yet optimised (Ferré et al., 2020). IVF in the bovine species is of particular interest, with its commercial scale now comparable to human IVF (Sirard, 2018). Likewise, for the equine, however intra-cytoplasmic sperm injection (ICSI) is currently the only commercial method for producing equine embryos *in vitro*. Research in the equine has recently focused on the early secretions of both the embryo and the maternal host (Swegen et al., 2017, Lawson et al., 2018, Smits et al., 2018, Fernández-Hernández et al., 2021) and highlighted the need for a more profound study of the oviduct. As species-specific differences between mammals are notable and varied, the traditional medical-model; beginning on laboratory rodents before clinical translation to humans (Bourdon et al., 2021) or mammal; is less than ideal. On these grounds, individualised organoid models derived from farm and companion animals have great potential to contribute to veterinary medicine as well as the health of the human population. The insights gained may benefit human fertility treatments and wildlife conservation efforts.

This study has succeeded in defining and characterising the growth and differentiation of oviductal organoids in domestic species and, in so doing, has demonstrated the effectiveness of a methodology that can be applied to future research on the reproductive biology of key livestock species. The development of an oviductal organoid model for domestic animal species allows direct access to the environment in which conception and early embryonic development occurs. As such it creates a platform for investigating the key attributes of a biologically supportive oviductal environment, which may include, but is not limited to further investigations into oviductal secretion, oviductal biomechanics, and the influence of the oviductal milieu on both embryo and spermatozoa. Such models will play a crucial role in addressing key issues relating to the efficacy of ART procedures, advances in animal welfare and the preservation of diversity. Furthermore, given that the oviduct is the site of sperm capacitation, fertilisation and early embryonic life, the findings of this study open up the potential to use this organoid model to achieve fundamental insights into the understanding of processes that are critical for the reproductive process both *in vivo* and *in vitro*.

## Supporting information

Svideo 1

## Author contribution

E.L., Z.G., R.J.A., P.T. Designed research; E.L., A.G., V.B., H.D. Performed research; H.D., R.L., M.B. Contributed new reagents; E.L., A.G., V.B., H.D. Analyzed data; E.L., M.B., Z.G., R.J.A., P.T. Wrote the paper.

## Acknowledgements

We would like to thanks members of the PRC for Reproduction, and the Australian Research Council, for supporting this work.

